# Structure-activity relationships of coumarin and its analogs and mechanistic insights into germination inhibition by coumarin

**DOI:** 10.1101/2024.05.23.595645

**Authors:** Kazuma Fukuda, Sota Hyakutake, Taiga Oishi, Michinari Yoshida, Mizuho Koga, Chisato Egami, Hyuga Matsuura, Ryusei Ito, Kosei Tsukahara, Mitsuki Noda, Kosei Tomonaga, Kohei Chifu, Rinsei Ichimaru, Eita Fujito, Kanae Mori, Yuka Tokuyama, Takako Yoshida, Noriko Ryuda, Yukio Nagano, Kazuhide Matsutaka

**Affiliations:** Saga Prefectural Chienkan High School, Saga, Japan; Analytical Research Center for Experimental Sciences, Saga University, Saga, Japan

## Abstract

Coumarin is produced by various land plants, such as cherry trees, and is known to exhibit a wide range of physiological activities. Notably, one of its primary functions is the inhibition of seed germination. However, the structural basis underlying coumarin’s germination-inhibitory effect remains poorly understood. In this study, we analyzed the structure-activity relationships of coumarin analogs in two plant species, white clover and Arabidopsis, and demonstrated that the benzene ring and ester bond within the coumarin skeleton constitute the minimal structural requirements for germination inhibition. The inhibitory effects of dihydrocoumarin, which lacks the double bond in the coumarin skeleton, varied among plant species, while those of various other modified coumarin analogs also differed across species. In both plant species, coumarin consistently proved to be one of the most potent inhibitors. Furthermore, the observation that dihydrocoumarin exhibited a strong germination inhibitory effect only in white clover suggests that certain coumarin analogs may act in a species-specific manner. Additionally, the mechanism of coumarin-mediated germination inhibition remains largely unclear. Using RNA-Seq analysis on Arabidopsis seeds, we uncovered a novel and critically important mechanism whereby coumarin suppresses the expression of genes encoding expansins, proteins that are responsible for loosening plant cell walls. This study is significant for understanding how coumarin contributes to plant competition in nature.

## Introduction

Coumarin, classified as a benzopyrone, is a heterocyclic compound composed of a benzene ring fused with a 2-pyrone ring that is produced by land plants[1,2]. In Japan, traditional products such as “sakurayu” (cherry blossom tea) and “sakuramochi”—a sweet made with mochi (Japanese rice cake) and preserved cherry leaves—derive their characteristic aroma primarily from coumarin [3]. Thus, coumarin is deeply intertwined with Japanese cultural heritage.

Coumarin is a versatile compound with a wide range of functions. In plants, it acts as an allelochemical by inhibiting the germination and root elongation of neighboring vegetation [4]. In this study, we focus on one of its primary roles: the inhibition of seed germination. The minimal structural elements required for coumarin-mediated germination inhibition remain unclear—that is, it is not known which parts of the coumarin skeleton are critical for this effect.

In addition, studies on the structure–activity relationships of coumarin-modified compounds are still limited.Numerous investigations have explored how modifications to coumarin affect seed germination. For example, using compounds such as 6-nitrocoumarin and 3-chlorocoumarin, Mayer et al. conducted a comparative analysis and found that structural modifications do not completely abolish coumarin’s germination inhibitory effect but only partially attenuate it [5]. This observation suggests that the ability to inhibit germination is conserved among coumarin analogs. Moreover, various coumarin analogs—including 4-hydroxycoumarin, 7-hydroxycoumarin (umbelliferone) [6], 4-methylumbelliferone [7], scopoletin (6-methoxy-7-hydroxycoumarin), Ayapin (presumed to be 6-acetoxy-7-hydroxycoumarin), and psoralen (a type of furanocoumarin) [8]—have been reported to exhibit germination inhibitory effects in different plant species. However, the relationship between molecular descriptors (such as molecular volume, molecular surface area, the lipophilicity index [LogP], and topological polar surface area [TPSA]) and the germination inhibitory effects of these compounds has not been thoroughly explored, highlighting the need for further research.

A critical aspect of understanding how coumarin influences seed germination is its effect on cellular processes. Several studies have investigated this mechanism, with the work by Chen et al. being particularly notable [9,10]. They demonstrated that coumarin negatively regulates the expression of abscisic acid (ABA) metabolic enzyme genes (OsABA8’ox2/3) and reduces levels of gibberellin A4 (GA4). Furthermore, coumarin suppresses the accumulation of reactive oxygen species (ROS) and the expression of key metabolic enzymes such as superoxide dismutase and catalase, thereby providing physiological evidence for the delayed germination observed upon coumarin treatment.

Omics analyses have significantly contributed to elucidating the mechanisms underlying germination inhibition [11,12]. For instance, Araniti et al. performed metabolomic analyses, revealing that coumarin extensively alters the seed’s metabolic profile by impacting amino acid metabolism, the citric acid (TCA) cycle, and aminoacyl-tRNA biosynthesis [11]. A similar study by Zhang et al. yielded consistent findings. Additionally, Zhang et al. conducted transcriptomic analyses 72 hours after coumarin treatment, demonstrating that coumarin affects the expression of genes involved in amino acid metabolism, the TCA cycle, and protein translation [12]. Although transcriptomic analysis has been used to investigate the biosynthetic pathway of coumarin [13,14,15,16], this study remains the only one to examine its germination inhibitory effect, rendering it invaluable. However, because these analyses were conducted 72 hours after treatment—by which time multiple downstream responses have likely been initiated (given that germination typically occurs one to two days after water absorption)—many previous studies may not have clearly identified the primary targets of coumarin-induced germination inhibition or fully accounted for the associated cellular structural changes.

In this study, we addressed these unresolved questions. First, we examined which structural elements of coumarin are minimally required for germination inhibition. Next, using coumarin-modified compounds, we investigated the correlation between germination inhibitory effects and molecular descriptors predicted by chemoinformatics, such as molecular volume and polar surface area. Finally, to evaluate the early effects of coumarin on germination inhibition, we conducted a transcriptomic analysis 24 hours after the addition of coumarin.

## Materials and Methods

### Evaluation of the importance of coumarin substructures in germination inhibition

Coumarin (Tokyo Chemical Industry, Tokyo, Japan) was employed as a positive control. For the assessment of the benzene ring, α-pyrone (Sigma-Aldrich Japan, Tokyo, Japan) was utilized, while 3,4-dihydrocoumarin (Tokyo Chemical Industry, Tokyo, Japan) was used for the evaluation of the double bond, and β-tetralone (Sigma-Aldrich Japan, Tokyo, Japan) was used for the assessment of the ester bond. These compounds were diluted with water and prepared at a concentration of 0.10 mmol/L. Seeds of white clover (Trifolium repens, purchased from Takii, Kyoto, Japan) were sown onto cotton-lined petri dishes, after which the solutions were added. Ion-exchanged water, without the addition of compounds, served as the negative control. Subsequently, the petri dishes were incubated in darkness at 20°C for 48 hours in an incubator (A4201, IKUTA Sangyo Co, Ltd, Nagano, Japan). For the germination test using Arabidopsis thaliana seeds (ecotype Col-0, purchased from Inplanta innovations, Kanagawa, Japan), in order to achieve a sufficient germination rate in the control group, the seeds were pre-soaked in 1 mL of solution at 20°C for 24 hours. After this pre-treatment, the germination test was conducted in the same manner as that for the white clover seeds. The number of germinations per petri dish was counted, and the percentage of germinated seeds out of the total number sown was calculated as the germination rate.

### Correlation analysis between cheminformatics-estimated indices and germination inhibition effects

Among the coumarin-modified compounds, five compounds were prepared: coumarin (KANTO CHEMICAL CO., INC., Tokyo, Japan), 7-methoxy coumarin (Tokyo Chemical Industry, Tokyo, Japan), aesculetin (Apollo Scientific Ltd, Manchester, UK), psoralen (Tokyo Chemical Industry, Tokyo, Japan), and umbelliferone (FUJIFILM Wako Pure Chemical Co., Osaka, Japan). The germination rates were calculated in the same manner as described above. Various indices estimated by cheminformatics for each compound were calculated using the following methods. The Topological Polar Surface Area (TPSA) and the octanol/water partition coefficient (LogP) were computed using SwissADME (http://www.swissadme.ch/) [17]. The molecular surface area and van der Waals volume were determined using winmostar (V11.5.8, https://winmostar.com/en/). The correlations between these indices and the germination rates were investigated. This analysis was performed on the germination rates of both Arabidopsis thaliana seeds and white clover seeds.

### Transcriptome analysis by RNA-Seq

Arabidopsis seeds, known for their well-characterized genetic information, were used. Initially, 20 mg of seeds were treated with either a 10 mmol/L coumarin solution or water and incubated for 24 hours. Compared to the structure–activity relationship experiments—which used 0.10 mmol/L due to solubility limitations—a higher concentration of 10 mmol/L coumarin was employed in the transcriptome analysis. This was because solubility was not an issue in this experiment, and using a higher concentration allowed us to clearly demonstrate its impact on the transcriptome. Three biological replicates were used for each condition. The seeds were then manually crushed with a pestle, and RNA extraction was performed using ISOSPIN Plant RNA (NIPPON GENE, Tokyo, Japan). Subsequently, RNA concentration was measured using the Qubit RNA BR Assay Kit (Thermo Fisher Scientific, MA, USA). RNA samples were sent to Novogene (Singapore) for quality testing using a Bioanalyzer (Agilent, California, USA), library construction using the NEBNext Ultra II RNA Library Prep Kit for Illumina, and sequencing on the NovaSeq 6000 platform (Illumina, California, USA). The resulting FASTQ data were analyzed using RaNA-seq (https://ranaseq.eu/) [18], where differential gene expression analysis was conducted based on the DESeq2 method [19]. Using the DESeq2 method with a pValue cutoff of 0.05 and a parametric Wald test, differentially expressed genes were detected. Genes with an adjusted p-value (padj) below 0.05 were considered significantly differentially expressed. PCA plots and MA plots were also generated using RaNA-seq. In addition, GSEA [20,21] was performed on these DEGs. The GSEA, including pathway and gene ontology analyses, determined the extent of impact by the normalized enrichment score (NES). The pathway analysis referenced the KEGG PATHWAY Database, and the gene ontology analysis was carried out with reference to the National Center for Biotechnology Information (NIH) database, analyzing biological process, cellular component, and molecular function domains.

### Real-time RT-PCR Analysis

The time-course analysis of expansin gene expression after coumarin treatment was performed using real-time RT-PCR. Arabidopsis seeds were treated with either a 10 mmol/L coumarin solution or water and incubated for 6, 12, 24, 36, and 48 hours, after which RNA extraction and quantification were performed as described above. cDNA synthesis was performed using the ReverTra Ace qPCR RT Master Mix with gDNA Remover (TOYOBO, Osaka, Japan) according to the manufacturer’s protocol. Real-time RT-PCR was carried out with the QuantStudio 3 Real-Time PCR system (Thermo Fisher Scientific, MA, USA) using the PowerUp SYBR Green Master Mix (Thermo Fisher Scientific, MA, USA) in triplicate. The primers used for real-time RT-PCR were as follows: ATEXPA1-F: 5’-TGCACGAGGCCTTGATGATG-3’, ATEXPA1-R: 5’-AACTCTCCCTCTCGCTTCGA-3’, EXPA2-F: 5’-ATGCTACCTTCTATGGTGGAGCTG-3’, EXPA2-R: 5’-AAAGCAGGCCCCACATTTCTG-3’, ATEXPA10-F: 5’-CCGTTAAAAAGGGCAAGTTGGTTAATCTCT-3’, ATEXPA10-R: 5’-GATTAGACGTCATCAAGGCCTTCTGCAAAA-3’, PP2A-F: 5’-ATCGCTACTCAGTTTAACCACA-3’, PP2A-R: 5’-TAGCAATAGTTTGGGGCACTAA-3’. The PCR program was as follows: 50°C for 2 min, 95°C for 2 min, followed by 40 cycles of 95°C for 1 s and 60°C for 30 s. The results were normalized to the expression level of PP2A, and the fold change relative to the 0 h time point was calculated.

### Utilization of generative AI

ChatGPT (https://chat.openai.com/) was used to rewrite the manuscript into more appropriate English and to proofread the text. The correctness of the revisions was subsequently verified by the authors.

### Ethics Statement

Permissions or licenses were not required for the laboratory use of these seeds. Additionally, there are no specific guidelines or legislation required for the use of the seeds in our study.

## Results

### Structural elements of coumarin minimally required for germination inhibition

Our experimental investigation into the minimal structural elements of coumarin required for germination inhibition revealed that the ester bond and benzene ring within its substructure are important for this activity, while the role of the double bond is species-dependent. In this study, we evaluated the germination effects on both white clover and Arabidopsis (S1 Table and S2 Table) seeds using compounds with substructures analogous to that of coumarin (Fig 1A): α-pyrone (Fig 1B), which lacks the benzene ring present in coumarin; β-tetralone (Fig 1C), which is missing both the double bond and the ester bond from coumarin’s lactone ring; and 3,4-dihydrocoumarin (Fig 1D), which lacks only the double bond from the lactone ring.

**Fig 1.**
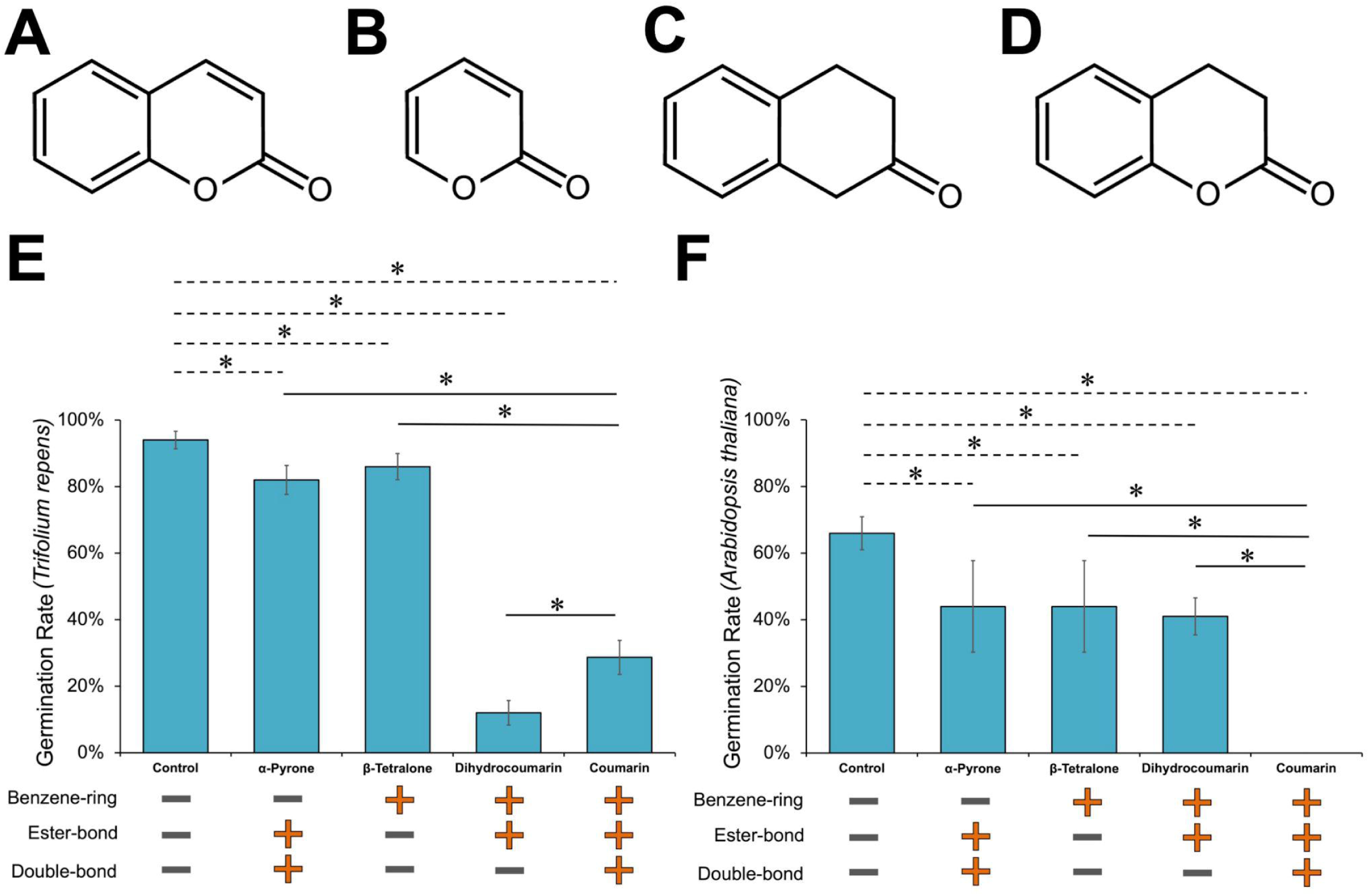
Effects of structural changes in the coumarin skeleton on seed germination. (A–D) Chemical structures of the coumarin-like substances used in this study are shown: coumarin (A), α-pyrone (B), β-tetralone (C), and 3,4-dihydrocoumarin (D). The solutions used in the experiments were prepared at a concentration of 0.10 mmol/L for both plant species. (E–F) Germination rates after 48 hours of incubation with the addition of coumarin-like substances are presented: (E) white clover (Trifolium repens), n = 300; (F) Arabidopsis thaliana, n = 300 for the control, 3,4-dihydrocoumarin, and coumarin, and n = 50 for α-pyrone and β-tetralone. Below the graph, the presence (+) or absence (–) of the benzene ring, ester bond, and double bond within the lactone ring for each substance is indicated. Group comparisons of germination rates were performed using chi-squared tests, with an asterisk (*) indicating statistical significance (p < 0.05). Error bars represent the 95% confidence intervals of the proportions. (Image source: Fig_1_0328.pdf)

For clover seeds, the germination rate for the negative control (water only) was 94% compared to 29% for coumarin-treated seeds. Seeds treated with α-pyrone and β-tetralone showed germination rates of 82% and 86%, respectively—values significantly higher than those of the coumarin-treated group and only slightly lower than the negative control (Fig 1E). In contrast, the germination rate for seeds treated with 3,4-dihydrocoumarin was 12%, markedly lower than both the coumarin-treated seeds and the negative control.

For Arabidopsis seeds, the negative control yielded a germination rate of 66%, while no germination occurred in the coumarin-treated group. Seeds treated with α-pyrone and β-tetralone exhibited germination rates of 44% and 44%, respectively—again, significantly higher than those for coumarin-treated seeds and only marginally lower than the control (Fig 1F). Notably, the germination rate for seeds treated with 3,4-dihydrocoumarin was 41%; although this rate was significantly lower than the negative control, it was considerably higher than that observed for coumarin, indicating a limited inhibitory effect.

Collectively, these studies using two different plant species demonstrate that the ester bond and benzene ring are important for coumarin-mediated germination inhibition, while the contribution of the double bond varies between species.

### Effect of coumarin-modified compounds on germination inhibition

After determining which substructures of the coumarin skeleton are critical for germination inhibition, we next examined how modifications to coumarin affect its inhibitory activity. Specifically, we focused on compounds that retain the important ester bond and benzene ring while preserving the double bond, given that the importance of the double bond for germination inhibition varies among plant species. Using these coumarin-modified compounds, we evaluated their structure–activity relationships.

Our results demonstrate that the inhibitory effects of modified coumarin analogs differ among plant species; however, coumarin consistently proves to be one of the most potent. To evaluate the factors influencing this activity, we investigated various molecular descriptors for the coumarin-modified compounds. TPSA is defined as the sum of the surface areas of all polar atoms—primarily oxygen and nitrogen, along with their attached hydrogens—in a molecule, reflecting its potential to interact with aqueous environments and traverse cell membranes. In addition, we calculated the molecular volume, which represents the three-dimensional space occupied by the molecule; the molecular surface area, defined as the total surface area of its three-dimensional structure; and the lipophilicity index LogP, defined as the logarithm of the octanol–water partition coefficient, serving as an indicator of hydrophobicity. These descriptors were determined for coumarin and the compounds shown in Fig 2A–D (S3 Table), and we examined the correlations between these values and germination inhibition.

**Fig 2.**
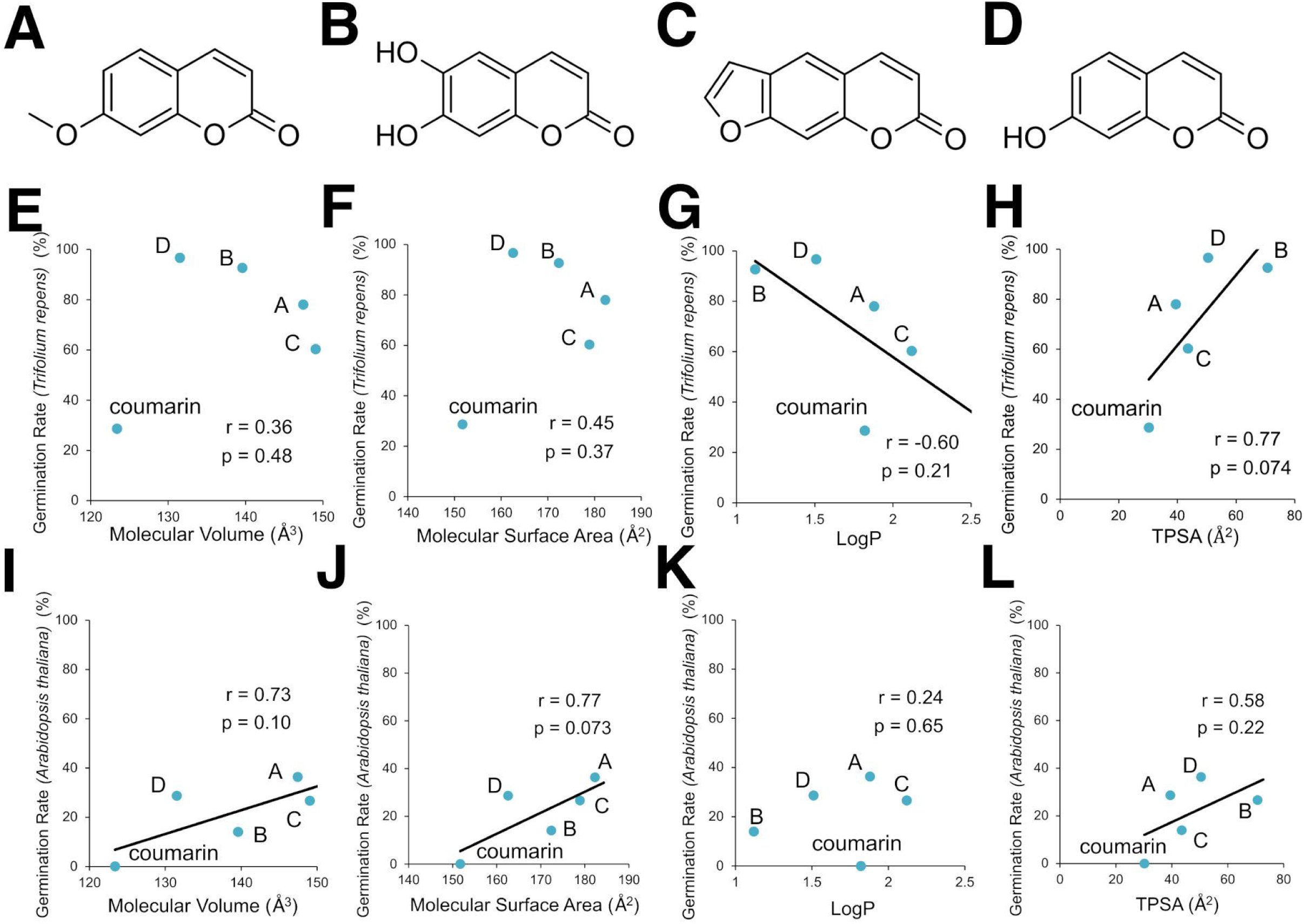
Effects of coumarin-modified compounds on seed germination. (A–D) Chemical structures of additional compounds used in this study, alongside coumarin as shown in Fig 1, are displayed: 7-methoxy-coumarin (A), aesculetin (B), psoralen (C), and umbelliferone (D). The solutions for the experiments were prepared at a concentration of 0.10 mmol/L for both plant species. (E–L) Relationships between the germination rates of seeds treated with the coumarin-modified compounds and four calculated structural indicators are shown (white clover (E–H) and Arabidopsis thaliana (I–L)). The y-axis represents the germination rates of seeds treated with each coumarin-modified compound (n = 300), and the x-axis represents the molecular descriptors calculated from their structures: Molecular Volume (E and I), Molecular Surface Area (F and J), LogP (G and K), and TPSA (H and L). Pearson’s correlation coefficient (r) and the p-value from the test of no correlation are provided. For each pair with an absolute correlation coefficient greater than 0.5, a scatter plot with a least-squares linear regression model is shown. (Image source: Fig_2_0328.pdf)

Based on a correlation coefficient threshold of r = 0.5, experiments using white clover seeds showed that a correlation was observed between germination inhibition and logP (r = –0.60) as well as TPSA (r = 0.77), but no correlation was found with molecular volume (r = 0.36) or molecular surface area (r = 0.45) (Fig 2E–H). Furthermore, in experiments using Arabidopsis seeds, correlations were observed between germination inhibition and molecular volume (r = 0.73), molecular surface area (r = 0.77), and TPSA (r = 0.58), but no correlation was found with logP (r = 0.23) (Fig 2I–L). In all experiments, the p-values calculated from the test of no correlation exceeded the commonly used threshold value of 0.05.

Thus, in both plant species, TPSA was the only molecular descriptor that showed a correlation with germination. Although this suggests that TPSA might be important, it should be noted that none of the p-values were sufficiently low. The key finding is that, although the effects of modified coumarin analogs vary among plant species, the compound that exhibited the strongest germination inhibitory activity is coumarin itself.

### Coumarin suppresses the expression of genes encoding expansin proteins

Utilizing Arabidopsis—a model organism with a well-established genetic profile—we examined the impact of coumarin on seeds using RNA-Seq analysis. Our results revealed that coumarin notably alters the expression of genes encoding expansins, which are responsible for loosening plant cell walls. The RNA-Seq analysis compared gene expression in seeds treated with coumarin for 24 hours to that in control seeds treated with water only. Three biological replicates were obtained for each group, yielding a total of six samples with an average of 3.9 Gbp of nucleotide sequences per sample. Principal component analysis (S1 FigA) demonstrated that the samples clustered into two groups (coumarin-treated and water-treated), with the coumarin-treated group exhibiting greater intra-group variability compared to the water-treated group. Using an adjusted p-value threshold of <0.05, we identified 3,416 differentially expressed genes (DEGs) out of 21,133 detected genes (S1 File). Differences in gene expression were shown in an MA plot (S1 FigB). Analysis of the DEGs revealed that many members of the expansin family (EXPA2/3/9/15, ATEXPA1/10) exhibited a significant reduction in expression (Fig 3A, S4 Table). Among these expansin genes, we further investigated the temporal expression of EXPA2, ATEXPA1, and ATEXPA10—which showed the most pronounced changes in RNA-Seq analysis—using real-time RT-PCR (Fig 3B–D). In the water-treated control group, the expression of these genes increased until 36 hours after water absorption and then declined. In contrast, in the coumarin-treated group, the expression levels were markedly suppressed. Based on the association with expansin genes, we performed a Gene Ontology (GO) analysis focusing on categories such as “cell wall,” “plant-type cell wall,” and “unidimensional cell growth” (S5 Table). This analysis revealed the enrichment of numerous expansin family genes within these categories. Additionally, other genes involved in xyloglucan metabolism, such as α-Xylosidase1 (XYL1) and members of the Xyloglucan endotransglucosylase/hydrolase (XTH) family, were also identified, with their expression data presented in Fig 3E (S6 Table and S7 Table).

**Fig 3.**
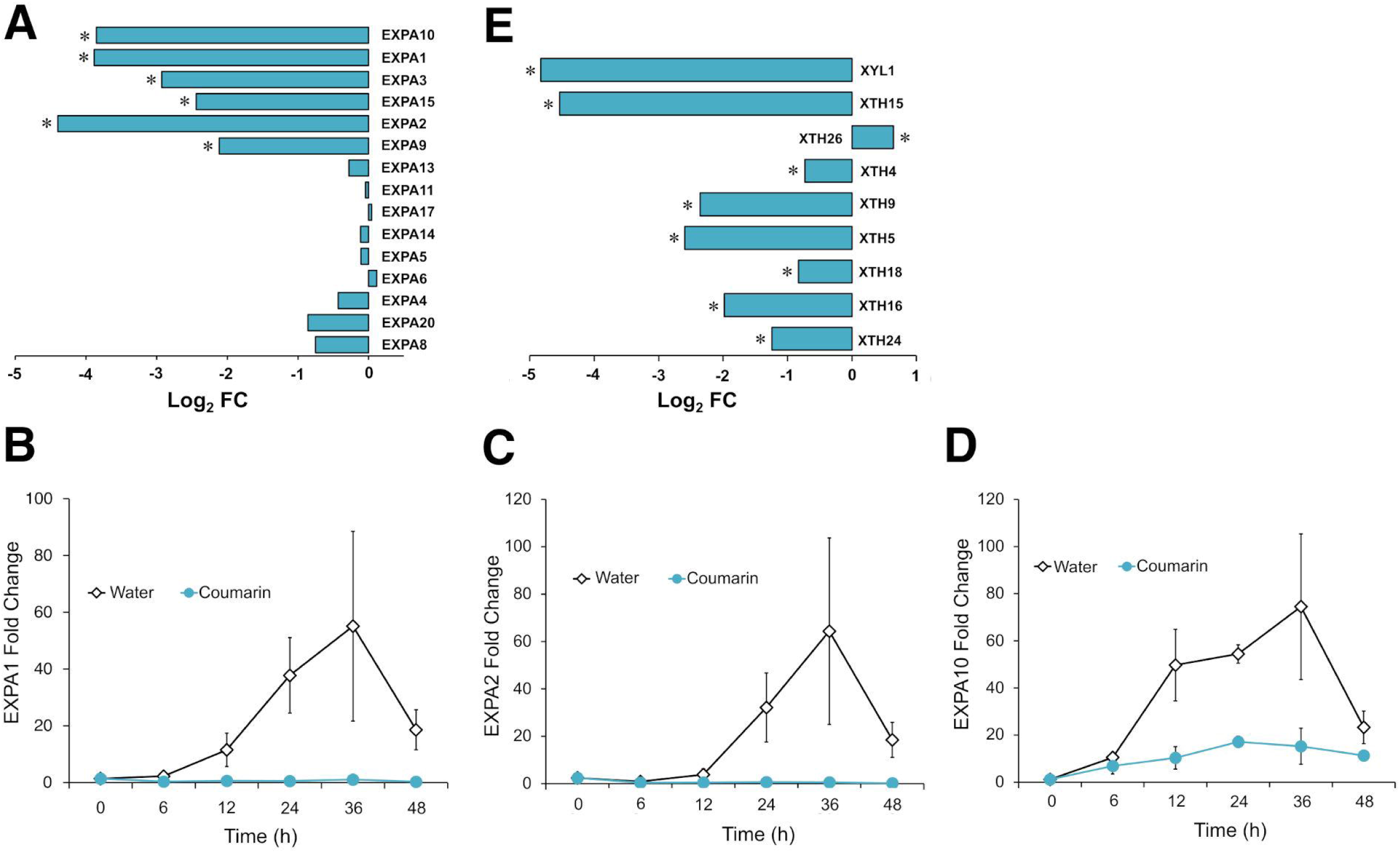
Down-regulation of expansin and cell wall-related genes by coumarin. (A) The bar plot shows the log2-transformed expression ratios of expansin family genes between coumarin-treated and water-treated control groups, as determined by RNA-Seq. (B–D) Real-time RT-PCR analysis of expansin family transcript levels: ATEXPA1 (B), EXPA2 (C), and ATEXPA10 (D). The data were normalized to the expression of the control gene (the PP2A gene encoding Protein Phosphatase 2A) and are presented as the fold change relative to the expression level at 0 hours. (E) Bar plots display the log2 expression ratios between the coumarin-treated and water-treated control groups for the XYL1 gene and members of the XTH family. An asterisk (*) indicates statistical significance based on the adjusted p-value (padj < 0.05). (Image source: Fig_3_0328.pdf) Note: The description for panel A in the original legend “Experiments were performed in triplicate for each group, with each column representing the expression profile of an individual sample” seems to describe a heatmap or multi-sample bar plot rather than the aggregated bar plot usually shown for log2FC. If panel A is an aggregated plot from RNA-Seq, this part of the legend might need adjustment. Assuming the plot shows mean log2FC values.

### Coumarin influences the expression of various genes

We utilized gene set enrichment analysis (GSEA) (Table 1) to investigate the impact of coumarin on gene expression. In coumarin-treated seeds, pathways related to ribosome function—specifically “Ribosome” and “Ribosome, eukaryotes”—were downregulated. Notably, genes encoding components of the large ribosomal subunit (RPL; 111 genes) and the small ribosomal subunit (RPS; 80 genes) were markedly suppressed (Fig 4A and 4B, S2 File and S3 File). Conversely, we observed upregulation in three pathways: “glutathione-mediated detoxification II,” “β-Oxidation,” and “Jasmonic acid biosynthesis” (S8 Table).

**Fig 4.**
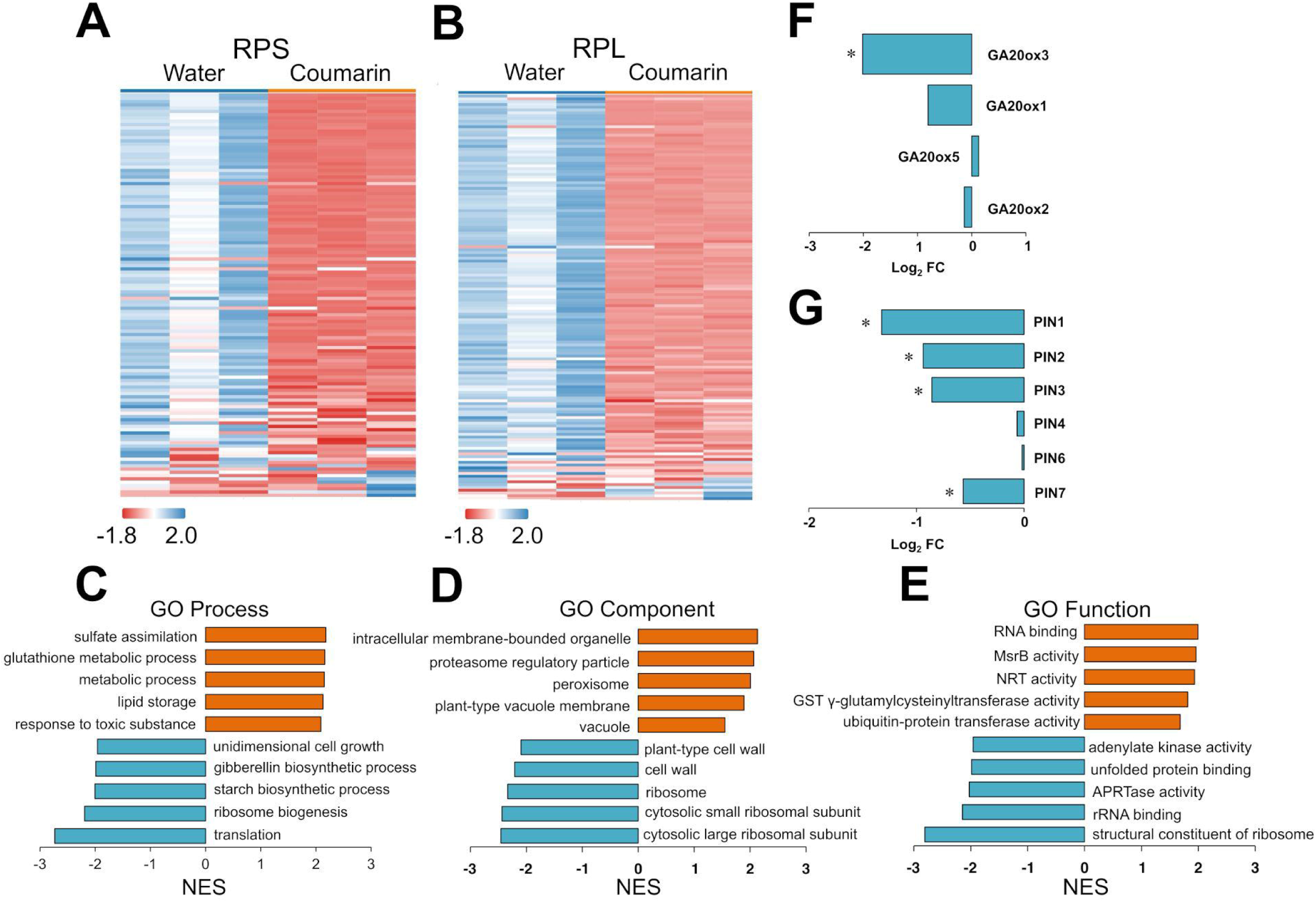
Coumarin Addition Impacts Multiple Pathways. (A–B) The heatmaps display the expression ratios of RPL family genes (A) and RPS family genes (A) between coumarin-treated and water-treated control groups as obtained by RNA-Seq. (C–E) Bar plots show the normalized enrichment scores (NES) from GSEA using the NIH Gene Ontology database. The top five activated and repressed pathways are presented for each gene ontology category: biological process (C), cellular component (D), and molecular function (E). (F–G) Bar plots depict the log2 expression ratios between the coumarin-treated and water-treated control groups for gibberellin 20-oxidase 1 and 3 (F) and for members of the PIN family (G). The asterisk (*) indicates statistical significance based on the adjusted p-value (padj < 0.05). (Image source: Fig_4_0328.pdf)

**Table 1.**
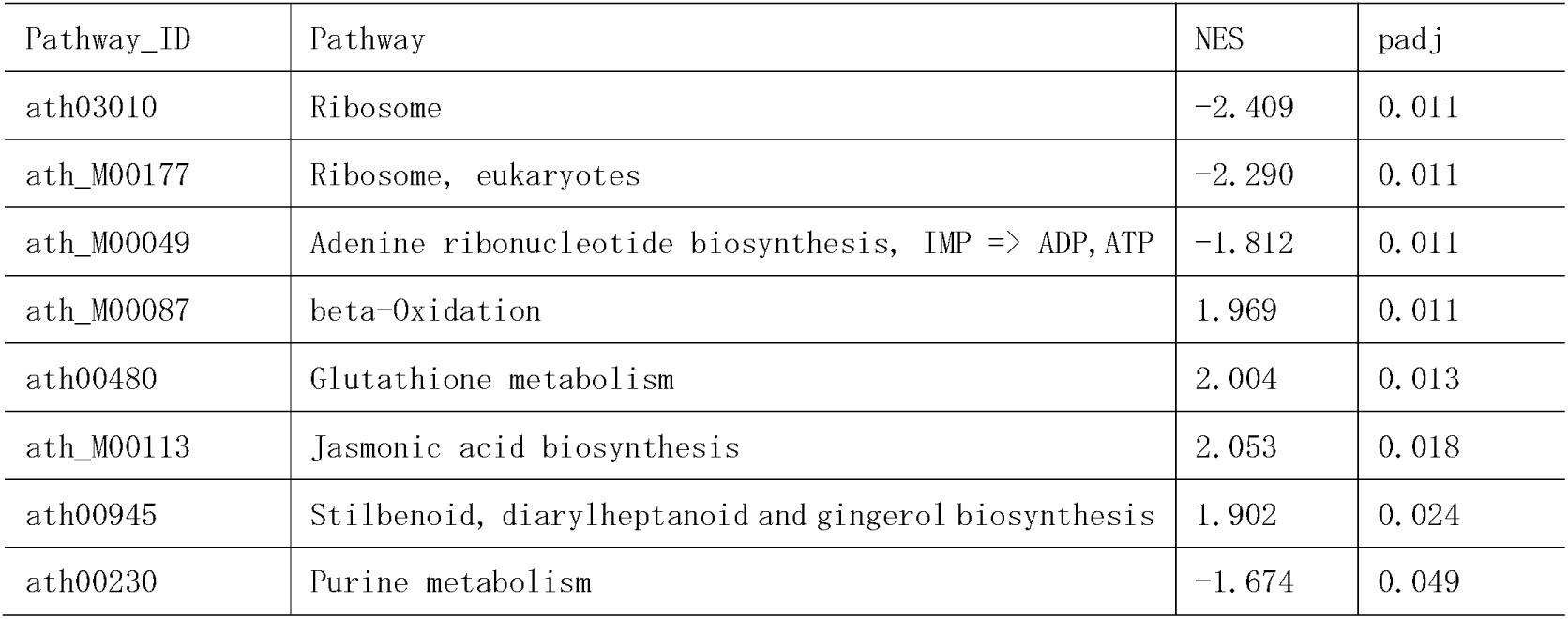
GSEA Pathway Analysis Results.

Further insights were obtained through Gene Ontology (GO) analysis (S4 File). The GO Process analysis revealed activation of genes involved in “sulfate assimilation” and “response to toxic substances,” whereas genes associated with “ribosome biogenesis,” “translation,” and “unidimensional cell growth” were inhibited (Fig 4C). Additionally, the GO Component analysis indicated that while genes encoding vacuolar components were upregulated, those forming part of the ribosomes and cell wall were downregulated (Fig 4D). Under the GO Function category, genes related to “RNA binding” and “glutathione gamma-glutamylcysteinyltransferase activity” were upregulated (Fig 4E).

Moreover, the expression of gibberellin 20-oxidase 1 (GA20ox1), a key enzyme in gibberellin synthesis, was significantly reduced (Fig 4F, S9 Table). Within the “unidimensional cell growth” category, PIN1—a member of the auxin transporter PIN family—and other PIN family members (PIN1/2/3/7) were also downregulated (Fig 4G, S10 Table).

## Discussion

One of the most significant findings of this study is that coumarin suppresses the expression of genes encoding expansins, which play a crucial role in loosening plant cell walls [22,23,24]. It is well established that the induction of expansin expression immediately following water uptake is critical for germination. For example, studies on tomato seeds have shown that LeEXP4 expression increases after water absorption, peaks at 24 hours, and then declines [25]. This pattern is consistent with our observation that EXPA2, ATEXPA1, and ATEXPA10 expression peaks at 36 hours after water uptake and subsequently decreases. Moreover, in the tomato study, puncture force analysis revealed that the weakening of the endosperm cap gradually progresses after water uptake, reaching a minimum at 36 hours in tandem with the expression pattern of LeEXP4, thereby supporting its role in germination 20. Additionally, research on Arabidopsis EXPA2 (EXP2) has demonstrated that its expression peaks at 24 hours after water uptake before declining [26]. Although there is a slight time lag—likely due to experimental conditions—the overall pattern of an initial increase followed by a decrease aligns with our observations. In that study, the delayed germination observed in an expa2 mutant further underscored the essential role of EXPA2 in germination. In summary, based on these previous studies and our findings, it is evident that one of the final targets of coumarin’s germination inhibitory effect is the loosening of cell walls mediated by expansins.

In the transcriptome analysis of Eleusine indica conducted by Zhang et al. [12], no suppression of expansin gene expression by coumarin was observed. In contrast, our analysis—performed 24 hours after water uptake—revealed significant suppression. This discrepancy is likely due to the timing of the analyses, as expansin gene expression is induced between 24 and 36 hours after water uptake and then declines. By analyzing gene expression at this earlier stage, we were able to detect the suppression of expansin gene expression by coumarin.

Our study indicates that the primary mechanism underlying coumarin-induced germination inhibition is the suppression of the expression of expansin genes, which are critical for loosening plant cell walls. Moreover, coumarin not only suppresses expansin gene expression but also downregulates genes involved in cell wall synthesis. In particular, we observed that the expression of genes encoding endoxyloglucan transferase A4 (XTH5) and α-xylosidase 1 (XYL1)—enzymes that interact with xyloglucan, a hemicellulose abundantly present in cell walls and essential for cell elongation [27,28]—is dramatically reduced by coumarin treatment. Collectively, these results suggest that coumarin exerts a multifaceted effect on cell walls to inhibit germination; further research is warranted to fully elucidate the underlying mechanisms.

Our research demonstrated that coumarin downregulates genes involved in the gibberellin synthesis pathway, linking this suppression to a decrease in cell wall loosening—a process critical for seed germination. Gibberellins (GA) regulate various germination-associated biological processes [29]. In addition, numerous studies have shown that GA induces expansin expression [30,31,32,33,34], with one particularly significant report confirming that GA induces expansin expression during seed germination 29. Notably, Chen’s discovery that coumarin reduces gibberellin synthesis further supports our transcriptomic findings [10].

It has been reported that coumarin interferes with polar auxin transport in the Arabidopsis root apical meristem [35]. Although the phenomenon we observed differs, we discovered that coumarin suppresses the expression of genes encoding members of the auxin transporter PIN family. In addition to its effect on the gibberellin signaling pathway, coumarin likely also influences the auxin signaling pathway.

A secondary discovery in this transcriptomic research is the significant suppression of genes encoding ribosomal proteins by coumarin. In line with our findings, Zhang et al. reported through transcriptomic analysis that coumarin also inhibits the expression of these genes in Eleusine indica [12]. Furthermore, their metabolomic analysis indicated that the synthesis of aminoacyl tRNAs is inhibited. While we examined gene expression 24 hours after water uptake and Zhang et al. conducted their analysis after 72 hours, both studies consistently show that coumarin suppresses the expression of genes encoding ribosomal components. Therefore, the translation system in seeds is likely another major target of coumarin. Given that ribosome biogenesis requires a substantial energy investment from the cell [36], it is conceivable that seeds exposed to chemical stressors like coumarin would suppress ribosome production. Moreover, since these suppressions were observed as early as 24 hours, it suggests that the inhibition of ribosome assembly occurs at an early stage, prior to the onset of additional cellular changes.

Furthermore, our study revealed that within the coumarin skeleton, the benzene ring and ester bond are important for its germination inhibitory activity. In contrast, the role of the double bond appears to be species-specific: it is non-essential in clover but plays an important role in Arabidopsis. Our Arabidopsis results are consistent with Mayer et al.’s findings, which indicated that the absence of the double bond weakens the germination inhibitory effect [5]. These observations suggest that the impact of dihydrocoumarin on germination is species-specific, reflecting differences in the underlying mechanisms of action.

Our research using two plant species suggests that the germination inhibitory effect of coumarin-modified compounds is correlated with their topological polar surface area (TPSA). In line with this observation, Mayer et al. demonstrated that the germination inhibition of modified coumarin compounds decreases in the following order: NO2 (TPSA: 49.36) > OH (TPSA: 20.23) > Cl (TPSA: 0) 5. These findings imply that higher TPSA values may enhance the compounds’ ability to interact with cellular targets, possibly by increasing cell membrane permeability or facilitating receptor binding [37]. However, our experiments with Arabidopsis revealed a weak correlation between TPSA and germination inhibition, suggesting that the influence of TPSA on activity may be modulated by species-specific physiological or biochemical factors.

Moreover, we did not observe significant correlations common to both plant species between germination inhibition and other molecular descriptors—namely, molecular volume, molecular surface area, and the lipophilicity index (LogP). Although these parameters are widely used in animal pharmacology and agrochemical research (e.g., in pesticide development), our results indicate that their impact on the inhibitory effect is limited.

An important observation is that while the inhibitory effects of various modified coumarin analogs differed among species, coumarin itself consistently proved to be one of the most potent germination inhibitors in both plant species examined. This consistency might be related to the ecological role of coumarin as a naturally produced compound in many plants, where its structure has been evolutionarily optimized for competitive interactions.

Despite the challenges in linking molecular descriptors directly to germination inhibition, the finding that dihydrocoumarin exhibited a stronger inhibitory effect than coumarin in white clover is particularly intriguing. This result suggests that specific modifications to the coumarin structure can enhance activity in a species-specific manner, opening up potential avenues for the development of targeted herbicides.

In summary, our study underscores the complexity of the structure–activity relationships in coumarin and its analogs. Future research should aim to identify additional molecular descriptors and investigate plant-specific factors that influence the bioactivity of these compounds. Such efforts will not only deepen our understanding of coumarin’s role in plant competition but also aid in the design of more effective, targeted herbicidal agents.

## Author Contributions

Conceptualization: K.F., T.O., S.H., M.Y., K.M.

Data curation: S.H., T.O., M.Y., H.M., M.K., C.E., K.T. (Kosei Tsukahara), K.T. (Kosei Tomonaga), K.C., R.Ich., E.F.

Formal analysis: S.H., T.O. (cheminformatics analysis), K.F., S.H., T.O., N.R., Y.N., K.M. (bioinformatics analysis), K.F., R.Ito (statistical analysis)

Investigation: S.H., T.O., M.Y., H.M., M.K., C.E., K.T. (Kosei Tsukahara), K.T. (Kosei Tomonaga), K.C., R.Ich., E.F. (germination tests), K.F., S.H., T.O., Y.T., K.M., T.Y., N.R. (RNA extraction), K.F., S.H., T.O., Y.T., K.M., T.Y. (real-time RT-PCR)

Methodology: K.F., T.O., S.H., M.Y., K.M.

Project administration: K.M.

Resources: N.R., Y.N.

Supervision: N.R., Y.N., K.M.

Validation: All authors.

Visualization: K.F., S.H., T.O.

Writing – original draft: K.F., S.H., T.O.

Writing – review & editing: Y.N., N.R., K.M., All authors.

## Supporting information

Supplementary Spread Sheets 1

Supplementary Spread Sheets 2

Supplementary Spread Sheets 3

Supplementary Spread Sheets 4

## Acknowledgments

The authors would like to express gratitude for the technical support provided by Nana Nakahara, Ayane Hirose, Itsuki Chiwata, Saori Nakagawa, Syunsuke Saeki, Yuki Takubo, Haruka Mine, Tatsuhiro Tanaka, Kotaro Yamamoto, Minoru Nishio, Masato Hara, Sota Fukushima, Shion Tanaka, and Hikari Kaneshige. RNA extraction and real-time RT-PCR experiments were facilitated at the Analytical Research Center for Experimental Sciences, Saga University. This work was supported by grants from the Nabeshima-Hokokai and the Japan Science and Technology Agency under the Super Science High School project of the Ministry of Education, Culture, Sports, Science and Technology.

## Data Availability Statement

The raw sequencing data for each sample have been deposited in the DDBJ Sequence Read Archive (https://www.ddbj.nig.ac.jp/dra/index-e.html; accession numbers DRR546110–DRR546115). All other relevant data are within the manuscript and its Supporting Information files.

## Supporting Information

S1 Fig. RNA-Seq quality control plots.

(A) Principal component analysis plot for RNA-Seq. This plot displays the samples along the first and second principal components. The orange and blue points represent the water-treated control group and the coumarin-treated group, respectively.

(B) MA plot of differential expression results. It shows the logL-scaled mean expression (x-axis) and the logL-scaled fold change (y-axis) for each gene in the differential expression analysis. Genes with significant changes in expression are highlighted as red dots.

S1 Table. Germination data for white clover with coumarin and analogs.

(This legend should briefly describe the content of S1 Table, which is referred to in source [48].

Example: “Germination rates and raw counts for Trifolium repens seeds treated with control, coumarin, α-pyrone, β-tetralone, and 3,4-dihydrocoumarin.”)

S2 Table. Germination data for Arabidopsis with coumarin and analogs.

(This legend should briefly describe the content of S2 Table, referred to in source [48]. Example: “Germination rates and raw counts for Arabidopsis thaliana seeds treated with control, coumarin, α-pyrone, β-tetralone, and 3,4-dihydrocoumarin.”)

S3 Table. Molecular descriptors for coumarin and modified compounds.

(This legend should briefly describe the content of S3 Table, referred to in source [68]. Example: “Calculated molecular volume, molecular surface area, LogP, and TPSA values for coumarin, 7-methoxy-coumarin, aesculetin, psoralen, and umbelliferone.”)

S4 Table. Expression data for expansin family genes from RNA-Seq.

(This legend should briefly describe the content of S4 Table, referred to in source [82]. Example: “Log2 fold change values and adjusted p-values for differentially expressed expansin family genes in Arabidopsis seeds treated with coumarin compared to water control.”)

S5 Table. Gene Ontology enrichment for expansin-related categories.

(This legend should briefly describe the content of S5 Table, referred to in source [86]. Example: “Enriched Gene Ontology terms (cell wall, plant-type cell wall, unidimensional cell growth) associated with expansin genes affected by coumarin treatment.”)

S6 Table. Expression data for the XYL1 gene from RNA-Seq.

(This legend should briefly describe the content of S6 Table, referred to in source [88]. Example: “Log2 fold change value and adjusted p-value for the XYL1 gene in Arabidopsis seeds treated with coumarin compared to water control.”)

S7 Table. Expression data for XTH family genes from RNA-Seq.

(This legend should briefly describe the content of S7 Table, referred to in source [88]. Example: “Log2 fold change values and adjusted p-values for differentially expressed Xyloglucan endotransglucosylase/hydrolase (XTH) family genes in Arabidopsis seeds treated with coumarin compared to water control.”)

S8 Table. Upregulated pathways from GSEA.

(This legend should briefly describe the content of S8 Table, referred to in source [92]. Example: “Details of significantly upregulated pathways (glutathione-mediated detoxification II, β-Oxidation, Jasmonic acid biosynthesis) from GSEA in coumarin-treated Arabidopsis seeds.”)

S9 Table. Expression data for gibberellin 20-oxidase genes from RNA-Seq. (This legend should briefly describe the content of S9 Table, referred to in source [97]. Example: “Log2 fold change values and adjusted p-values for GA20ox1 and GA20ox3 genes in Arabidopsis seeds treated with coumarin compared to water control.”)

S10 Table. Expression data for PIN family genes from RNA-Seq.

(This legend should briefly describe S10 Table, referred to in source [98]. Example: “Log2 fold change values and adjusted p-values for differentially expressed PIN family genes in Arabidopsis seeds treated with coumarin compared to water control.”)

S1 File. List of all differentially expressed genes (DEGs).

(This legend should briefly describe S1 File, referred to in source [80]. Example: “Complete list of 3,416 differentially expressed genes in Arabidopsis seeds treated with coumarin for 24 hours compared to water control, including log2 fold changes, p-values, and adjusted p-values.”)

S2 File. List of differentially expressed Large Ribosomal Subunit (RPL) genes.

(This legend should briefly describe S2 File, referred to in source [91]. Example: “List of 111 differentially expressed genes encoding components of the large ribosomal subunit (RPL) in coumarin-treated Arabidopsis seeds.”)

S3 File. List of differentially expressed Small Ribosomal Subunit (RPS) genes.

(This legend should briefly describe S3 File, referred to in source [91]. Example: “List of 80 differentially expressed genes encoding components of the small ribosomal subunit (RPS) in coumarin-treated Arabidopsis seeds.”)

S4 File. Complete Gene Ontology (GO) analysis results.

(This legend should briefly describe S4 File, referred to in source [93]. Example: “Detailed results of Gene Ontology (GO) enrichment analysis for biological process, cellular component, and molecular function categories for differentially expressed genes in coumarin-treated Arabidopsis seeds.”)

