## Supplementary figures and images for "Structure-activity relationships of coumarin and its analogs and mechanistic insights into germination inhibition by coumarin"

### Supplementary Figure 1

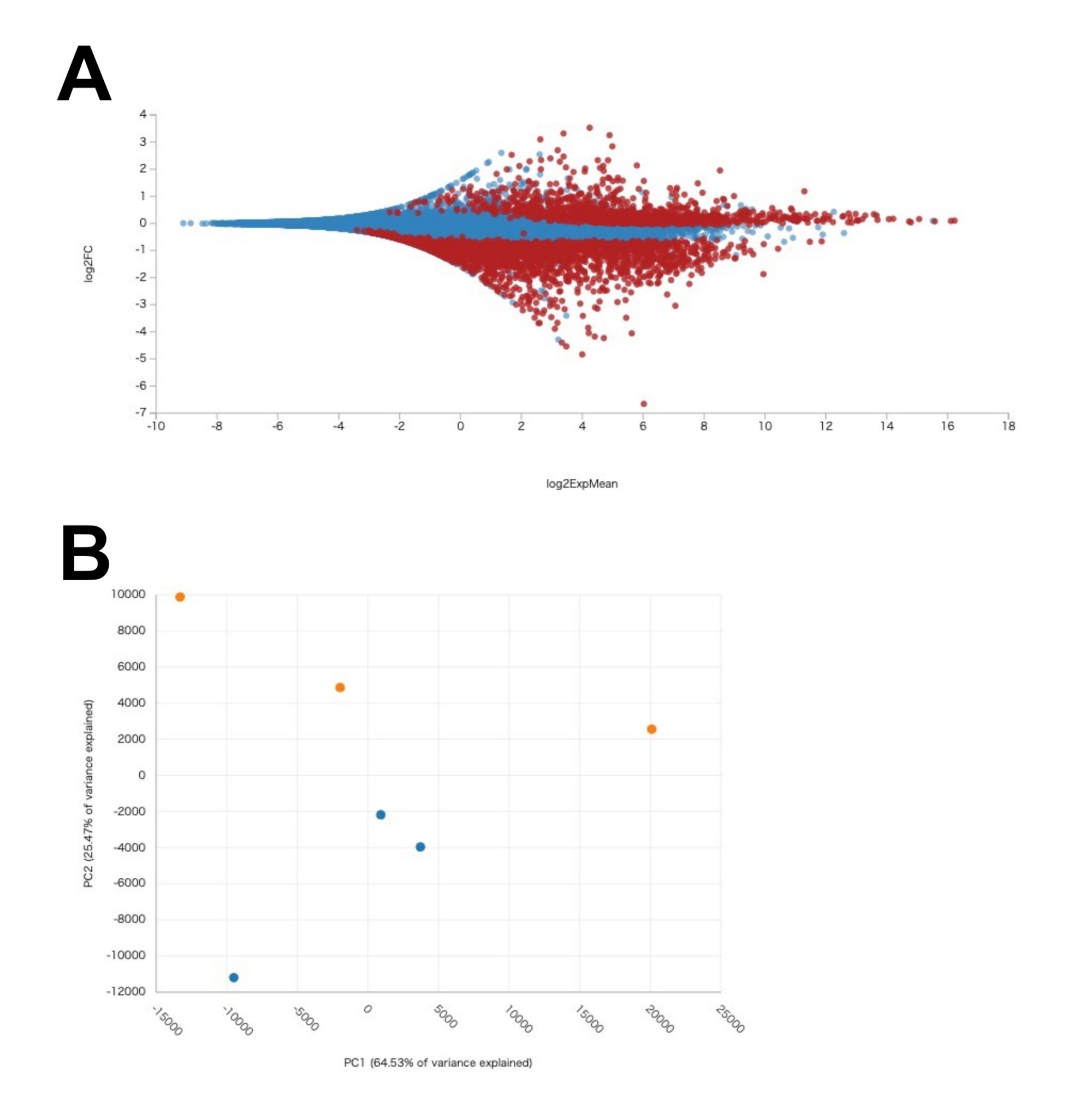
